# Spirals, Scalp Whorls and Skin Biomechanics: Nature’s Own Design For Expansion

**DOI:** 10.1101/043992

**Authors:** Sharad P. Paul

## Abstract

This paper began as an exercise in curiosity – logarithmic spiral designs abound in nature ‐‐ in galaxies, flowers, pinecones and on human scalps as whorls. Why are humans the only primates to have whorls on the scalp? Is the formation of scalp whorls mechanical or genetic? A mechanical theory has long been postulated– the mechanical theory suggests that hair whorl patterning is determined by the tension on the epidermis during rapid expansion of the cranium while the hair follicle is growing downwards – however, this has never before been experimentally proven conclusively. We found, that under certain conditions, we were able to experimentally recreate spirals on the scalp to demonstrate that the basis of scalp whorls is indeed mechanical – and that logarithmic spirals are indeed nature’s own design for rapid expansion of organic tissues. Given our experiments only created whorls when certain conditions were satisfied (and not in others), they have given us great insight into the mechanical formation of skin whorls and the physiology of skin stretch. We believe that these findings will lead to many more advances in understanding skin dynamics and indeed the behavior of any living tissue when confronted by stretch. As a corollary, the application of the results of these studies have led us to the discovery of a new surgical technique for closure of scalp defects using the golden spiral pattern, and this will be the subject of a separate paper.

## Introduction

Logarithmic spirals such as golden spirals (with a growth factor of φ = 1.6180339887, the golden ratio) are ubiquitous in nature and are seen in arrangements of leaves, seeds, pinecones and many different arrangements in nature.^1^ Such spiral patterns are frequently observed and utilized in a variety of phenomena including galaxies, biological organisms, as well as turbulent flows.^2^ Many have attributed mysticism to the presence of origins – the golden ratio was called τ (tau), the symbol of life, and was also known to ancient Egyptians (The ankh) and Hindus.^3^ One of the viewpoints for the occurrence in plants was inherently protoplasmic i.e. “spirals” exist in nature through constitutional rather than external mechanical influences.^4^ However, a team noted that metallic nanoparticles could be made to arrange themselves with floral spiral design and commented that the shape of spirals depended on size of particles, thickness of shell and rate of cooling.^5^ Others have stimulated spiral wave forms in neuronal circuits of the human body.^6^ While Fibonacci spirals are everywhere in nature, and indeed common in art and architecture ‐‐ in a laboratory setting, spontaneous assembly of such patterns has rarely been realized in mechanical tests, and never before on skin. ^6^ Some researchers demonstrated Fibonacci spiral patterns could be reproduced through stress manipulation on the metal-core based shell microstructures and commented on the possible role of mechanical stress in influencing these spiral patterns. ^7^ And, in a study of rocks and tectonic plates, researchers noted spiral patterns of cracks ‐‐ where the cracking arises not from twisting forces but from a *progressing stress front*.^8^

But as an animal biologist and cutaneous surgeon, my fascination has been with the fact that humans are the only animals to have scalp whorls on the top of their heads that follow spiral patterns seen in nature. Others have noted that of all mammals, only humans have hair whorls on the vertex of the scalp, and that each human individual *must* have a hair whorl.^9^ Animals like horses do have whorls on the body or faces, but not on the top of their heads – and this is not universal in all horses – however, such patterns demonstrate less variability, and show more Mendelian forms of inheritance than those seen in humans.^10^ In keeping with such genetic theories, the number of whorls on a horse’s face and the direction of such patterns are used to gauge equine temperament such as calmness, enthusiasm or wariness.^11^ Notably, other primates like chimpanzees, that are evolutionarily closer to humans, do not have whorls.^12^ In this experiment we re-created skin whorls by causing shearing forces due to rapid skin expansion along a progressing front.

## Methods

### The Medium

The author spent the past year in testing and studying the properties of pigskin, and its suitability as an equivalent and comparable testing medium to human skin, before formally testing the mechanical theory of spiral formations in organic tissue.

Before one can study skin bio-dynamics or surgical closure techniques, one needs a good understanding of the directions of skin tension lines of normal skin. Karl Langer (1819-1887), Professor of Surgery in Vienna, was one of the earliest people to undertake such studies, when he undertook experiments to study physical and mechanical properties of human skin^13^ in cadavers by making small circular incisions on skin – and then noting the directions they were distorted in – these ‘skin tension lines’ he termed as ‘cleavage lines.’^14^ Since Langer’s original study, many other variations of skin tension lines have been mapped out ‐‐ and people have noted that in certain parts of the face, Langer’s lines deviate from relaxed skin tension lines – nevertheless Langer’s work remains the most important in any discussion regarding skin tension lines.^15^ More recently, investigators studied porcine skin to assess the presence of ‘Langer’s Lines’ and concluded that, as in humans, Langer’s Lines do have specific patterns in pigskin^16^ and are equally dynamic with movement – with the authors noting that in pigskin the ‘tension lines were oblique to the vertebra in the cephalic area, perpendicular to the vertebra in the middle torso, and parallel to the vertebra in the caudal area.’^16^ One of the other factors I took note of prior to testing was the relationship between Langer’s Lines and hair follicles – experimental studies have shown that there is a good correlation between Langer’s Lines and direction of hair streams^16^ and this knowledge, hitherto helpful in studying wounds, also helped in the design of my experiment – especially in planning skin markings and incisions for this study.

The mechanical theory of scalp whorl formation suggests that during the 10^th^ to 12^th^ week of foetal life, the brain expansion is so rapid that it creates a shearing force between the two layers of skin, at the dermo-epidermal junction.^17^ While planning for this study, I conducted expansion studies using saline expanders to understand the pressure required to create such shearing forces. In a cross section of pigskin, as in Fig.1 one is able to observe the epidermis and a layer of adipose tissue that serves as a fatty ‘dermis’ layer. Using saline expanders at the margins of pigskin cross sections allowed us to observe dermo-epidermal shear. I found rapid expansion resulted in enough shearing force to separate the layers of skin. Slow expansion over days or weeks did not result in creating enough ‘shear’ in pigskin. Pigskin has three separate fat layers, separated by two layers of fascia.^18^ The uppermost layer of fat in pigskin corresponds to the dermis in human skin – and authors have coined the term ‘intradermal adipocytes’ to describe these cells, as the term accurately reflects both their developmental origin and anatomical location.^19^ Therefore, for the purposes of this experiment, I decided to see if the causation of shearing forces between the uppermost two layers of pigskin (the epidermis and fatty dermis) would result in the formation of spirals along an advancing front.

**Fig. 1.**
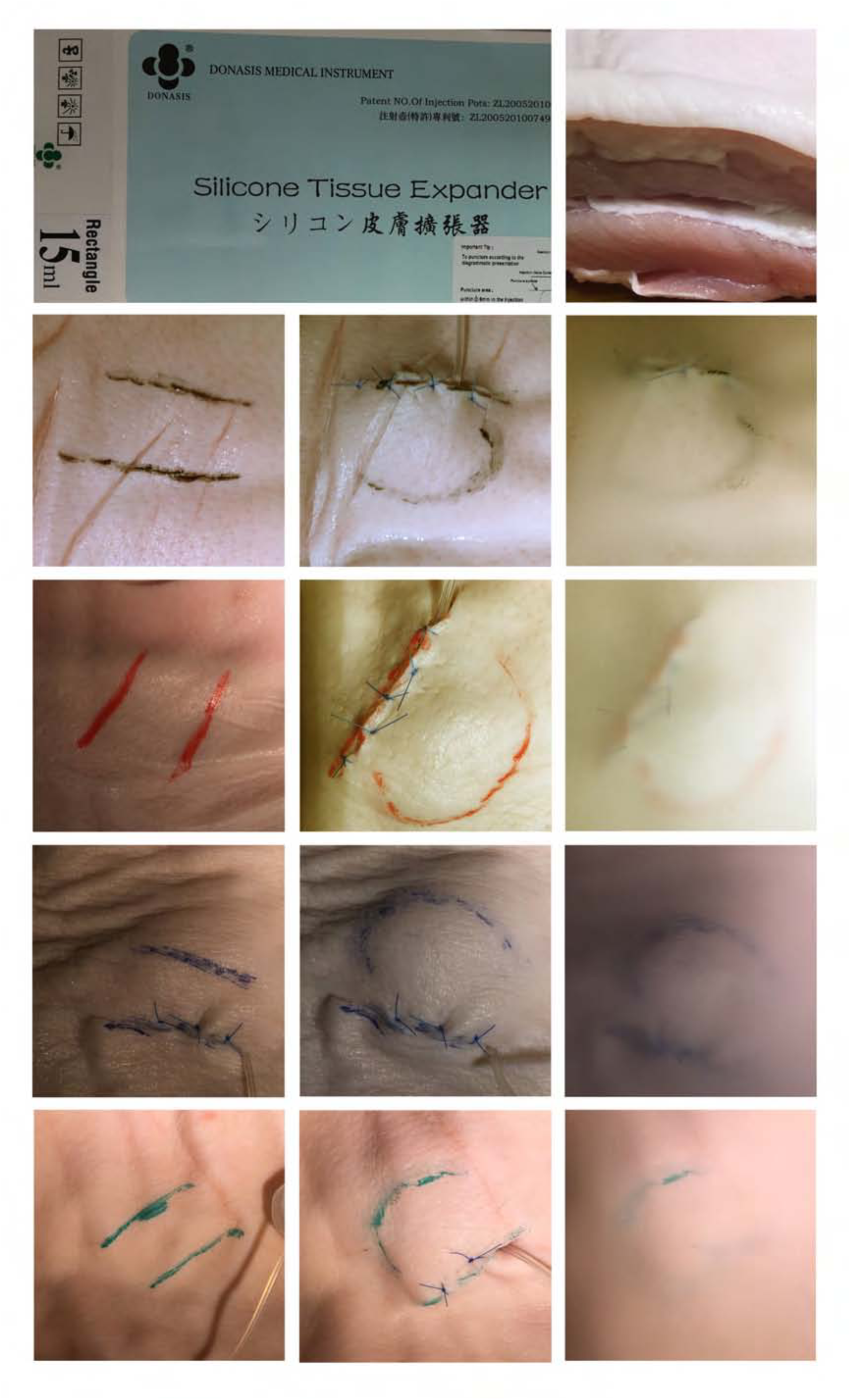
(Skin tension lines curving along an advancing front due to rapid saline expansion.)

### The Experiment

Twenty freshly slaughtered pig bellies were obtained from an abattoir. Two parallel skin lines were marked along hair-follicle directions on pigskin, thereby indicating two parallel skin tension lines, about 2 cm apart. These were marked along ‘Langer’s Lines’ on the pig and not along wrinkle lines (photographs show that in some cases, wrinkle lines ran across the skin tension lines). Permanent indelible ink markers were used for each of these markings. A 15 ml silicone tissue expander was used in each case (Shanghai Wushan Industrial Co., Ltd, under license from Donasis Bio Labs, Japan). I deliberately used a rectangular silicone expander to ensure that any spirals formed were not merely due to the shape of the implant used. A 2cm incision was made in the skin, and was deepened to the muscle layer below the ‘dermis’ in the pig. A pocket was created, only wide enough to accommodate the collapsed silicone expander – just wide enough to fit the silicone expander that was firmly compressed and squashed into position. The incision was closed in layers. Care was taken to incorporate deeper layers within the deeper sutures to make sure that the ‘advancing front’ due to this rapid tissue expansion would be in one direction only (away from the suture-line) i.e. one of the two tension lines marked would be fixed to underlying tissues and not capable of expansion. The expanders were filled with saline rapidly (for purposes of this experiment, from previous testing, we defined this as expansion of the silicone expander with 10 ml of saline in under 2 seconds, followed by a further expansion to full capacity after 2 minutes). Tracing paper was placed over the pigskin and the curvature of the advancing line due to tissue expansion was marked out. The process was identical in each case, except we used a different pig belly in each case, and markings were traced using different coloured inks.

The reason for the two-stage rapid expansion is because such staged protocols have been established in surgery for acute tissue expansion^20^, with many authors supporting this method of rapid expansion to achieve additional skin cover during reconstructive surgery.^22^ Further, during the testing phase, I had found this technique consistently caused enough dermo-epidermal shear to cause displacement of the fatty dermis. During such acute expansion of tissue, biological and mechanical ‘creep’ occurs, and authors have postulated that it is the displacement of water from the collagen network and micro-fragmentation of elastic fibers that makes skin more viscous.^22^ To mitigate any variability of technique and ensure accuracy, the same person (in this case the author) was the only person performing all the testing.

## Methods Discussion

In our experiment, skin tracings of the advancing front (i.e. the non-sutured skin tension line that becomes deformed into curves under the force of rapid expansion) were marked out as detailed above (Fig. 1). To the naked eye, while we had achieved some degree of displacement of the original straight-line markings into curves, they did not seem to confirm to any particular spiral pattern (Fig. 2). However, when we had traced out all the patterns and then superimposed all our tracings, one on top of each other to create a composite image, we noted a distinctive logarithmic spiral pattern. The result was both unexpected and astonishing – the composite image created by overlaying all our tracings show a clear logarithmic spiral pattern, in our case very close to a golden spiral pattern seen in nature. Fig. 2 shows each individual tracing, followed by a composite image, finally overlaid by a golden spiral. Each tracing, when analyzed retrospectively did indeed conform to a portion of a golden spiral.

**Fig. 2.**
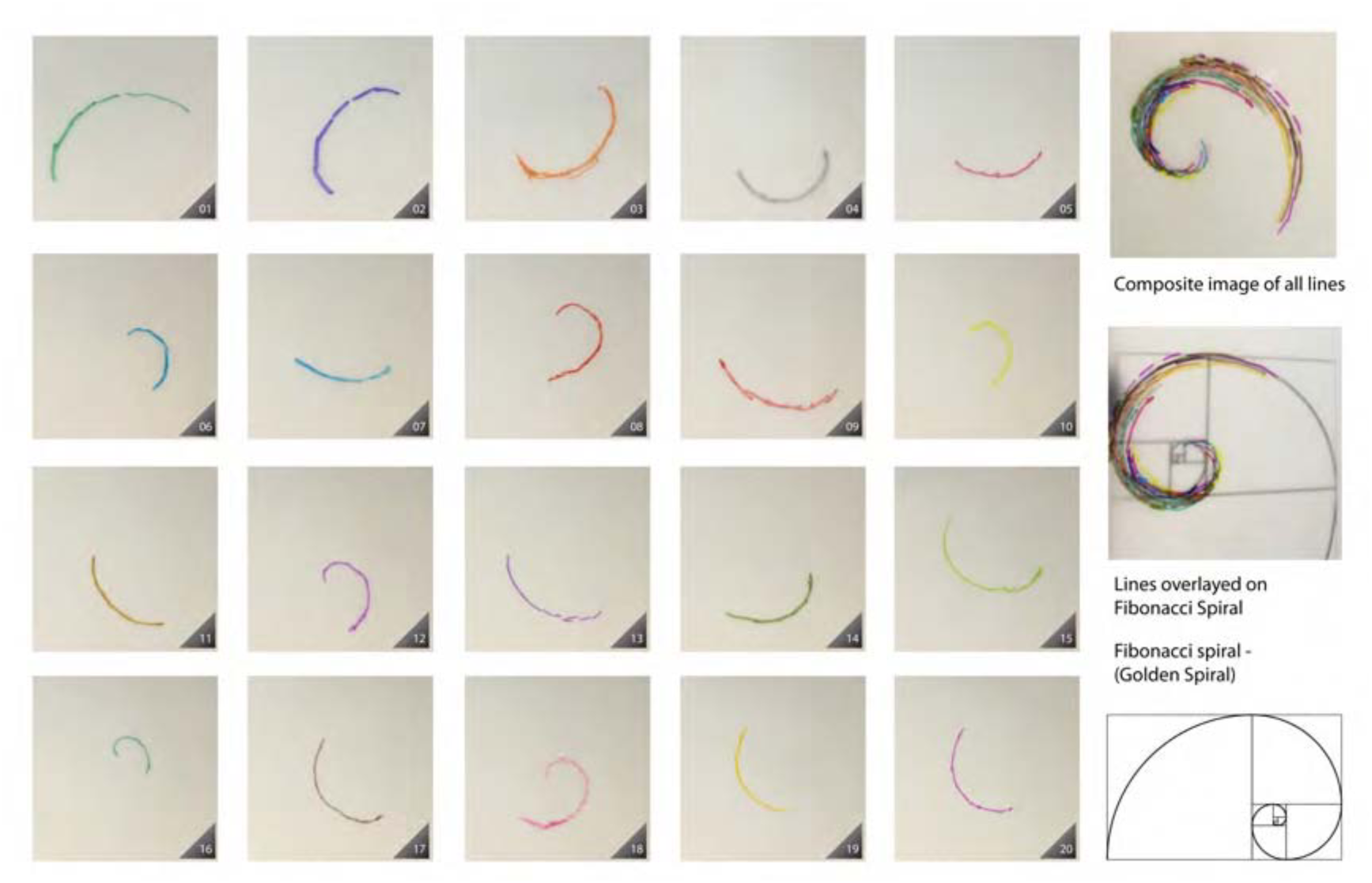
Skin tension lines marked out using tracing paper, followed by a composite image

This experiment demonstrates clearly that under certain conditions, rapid tissue expansion causes dermo-epidermal shear, which can result in the formation of spirals – in our case, nature’s own design: the golden spiral. Knowledge gained from this experimental study has helped the author design a new technique for closing scalp defects, and this will be the subject of a separate paper in a surgical journal.

As we discussed earlier, spiral formations abound in nature and in tissues, they may represent a feature of rapid expansion along an advancing front. After all, the opening of flowers is often rapid ‐for example, Hedera helix, the English Ivy, opens in about 5 minutes^23^, and many flowers demonstrate spiral patterns around their inflorescence axis. The opening of flowers is already known to be due to cellular expansion – while floral openings can be classified as nocturnal, diurnal, single or repetitive, previously published papers have demonstrated that the opening of flowers is generally due to cell expansion.^24^ As to the genetics of such rapid expansion, researchers have studied rapid intraoperative tissue expansion in mouse skin and found that in response to stretch caused by a balloon, similar to that used in our experiment, the following genes are induced ‐‐ L1 (truncated long interspersed nucleotide element 1), myotubularin and insulin.^25^ In another study on adult skin, researchers found a significant difference in 77 genes after expansion – with a significant finding being the expression of regeneration related genes, such as HOXA5, HOXB2 and AP1, after tissue expansion, with implications for further research into skin regeneration.^26^ Indeed studies in monozygotic twins have shown individual variations of scalp whorls, and therefore both skin stretch and the subsequent gene expression can be considered responsible for the formation of spirals.^27^

The mechanical theory of scalp expansion causing whorls, while hypothesized, has never been re-created in a controlled experiment on skin. This study therefore represents a substantial advance in the understanding of both the formation of scalp whorls and specifically, the golden spiral – which can be considered nature’s own designs for rapid expansion of organic matter, along an advancing front. Given this has the potential for far-reaching consequences, and further research in other fields of scientific endeavour, I am presenting this to the wider scientific community.

## Acknowledgements

I would like to thank Jan Gardner at Auckland University’s Advanced Clinical Skills Centre for helping obtain pigskin and use of the Centre’s facilities for my experiments. This paper resulted from a study that forms part of my Ph. D research project at the University of Queensland’s School of Medicine and I would like to acknowledge my supervisor, Assoc. Professor Cliff Rosendahl. And finally, I would like to thank Ryan Butler, Auckland University of Technology, for his help with photographs to document my findings.

